# Membrane oscillations driven by Arp2/3 constrict the intercellular bridge during neural stem cell divisions

**DOI:** 10.1101/2024.10.28.620743

**Authors:** Bryce LaFoya, Kenneth E. Prehoda

## Abstract

After the first furrowing step of animal cell division, the nascent sibling cells remain connected by a thin intercellular bridge (ICB). In isolated cells nascent siblings migrate away from each other to generate tension and constrict the ICB, but less is known about how cells complete cytokinesis when constrained within tissues. We examined the ICBs formed by *Drosophila* larval brain neural stem cell (NSC) asymmetric divisions and find that they rely on constriction focused at the central midbody region rather than the flanking arms of isolated cell ICBs. Super-resolution, full volume imaging revealed unexpected oscillatory waves in plasma membrane sheets surrounding the ICB pore during its formation and constriction. We find that these membrane dynamics are driven by Arp2/3-dependent branched actin networks. Inhibition of Arp2/3 complex activity blocks membrane oscillations and prevents ICB formation and constriction. Our results identify a previously unrecognized role for localized membrane oscillations in ICB function when cells cannot generate tension through migration.

## Introduction

Cell division in animal cells proceeds through three sequential membrane constriction steps^1^ (Fig. 1A). First, a broad actomyosin ring rapidly constricts the plasma membrane between the segregated chromosomes until the connecting pore reaches a diameter of ∼1.5 μm. The intercellular bridge (ICB) – a membranous tube containing the compacted central spindle – connects the nascent siblings after the initial furrowing step^2^. The ICB then undergoes a slower constriction phase, narrowing to ∼200 nm in diameter, before the final step of abscission severs the membrane connection between the daughter cells^3–5^. While the molecular mechanisms driving initial furrowing and final abscission are relatively well understood, the intermediate phase of ICB constriction remains poorly characterized. Most insights come from studies of cultured cells, where the ICB forms a long, thin tube as the nascent siblings actively migrate apart^2,6^. This ICB consists of a central midbody flanked by constricting arms (in an alternate nomenclature ICB and midbody are synonymous and the central structure is the stem body)^2,7–10^. When the arms narrow to ∼200 nm, the midbody-recruited ESCRT-III resolves the membrane during abscission^3,11–13^. Here we address how ICB constriction occurs in cells that are part of a tissue, where mechanical constraints may preclude the nascent sibling separation that occurs in cultured cells.

**Figure 1:**
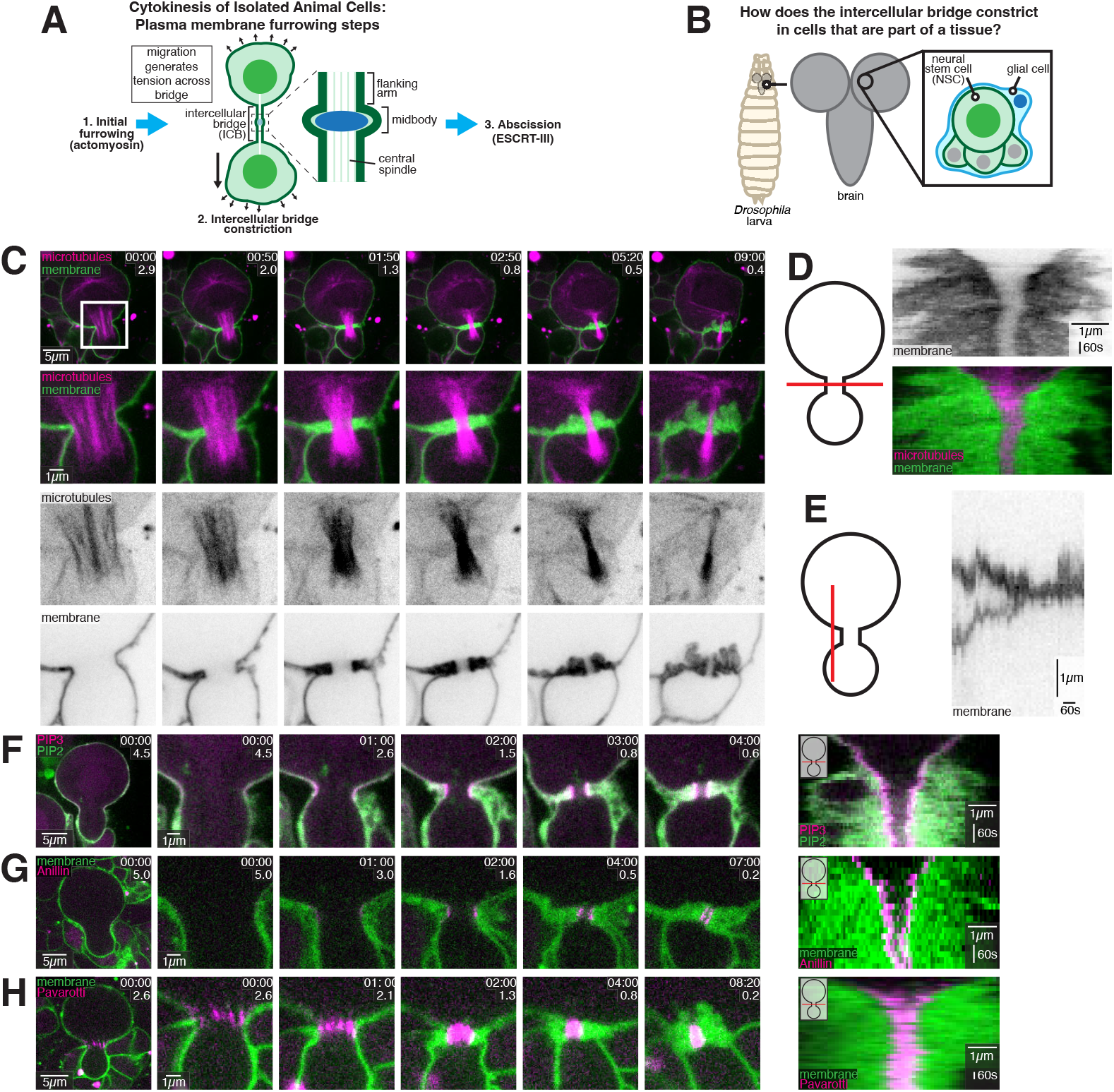
Larval brain neural stem cell intercellular bridges are midbodies without flanking arms. A) Intercellular bridge (ICB) constriction. Animal cell cytokinesis consists of three furrowing steps, the initial furrowing of the actomyosin ring, ICB constriction, and abscission. In isolated cells like cultured cells, the ICB is constricted by migration of the nascent siblings away from one another, which generates tension across the bridge. Constriction, or “thinning” of the ICB occurs at the flanking arms, leaving the central bulge from the midbody. B) Larval brain neural stem cells (NSCs) are part of a complex tissue. *Drosophil*a larval brain NSCs are surrounded by a cortex glial cell (blue) and progeny from previous divisions (smaller cells). This study uses the NSC as a model system for understanding how ICB constriction occurs in cells that are part of a tissue, where mechanical constraints may preclude the mechanisms used by isolated cells. C) Membrane and microtubule dynamics during the late stages of NSC division. Selected frames from Video 1 are shown. Time in minutes relative to the start of imaging is shown, along with the diameter of the cytokinetic pore in microns. D) Kymograph of membrane and microtubule dynamics across the cytokinetic furrow as the intercellular bridge forms and thins. E) Kymograph demonstrating that during cytokinesis, nascent sibling cells are initially distanced from one another and then come in contact. F) Phosphoinositide localization during the late stages of NSC division. Selected frames from Video 1 are shown. The NSC is expressing UAS-GRP1-GFP “PIP3” and UAS-PLCδ-PH-mCherry “PIP2”. Time in minutes relative to the start of imaging is shown, along with the diameter of the cytokinetic pore in microns. To the right is a kymograph of phosphoinositide localization across the cytokinetic pore. G) Anillin and membrane dynamics during the late stages of NSC division. Selected frames from Video 1 are shown. Time in minutes relative to the start of imaging is shown, along with the diameter of the cytokinetic pore in microns. To the right, a kymograph of Anillin and membrane dynamics across the cytokinetic pore is shown. H) Pavarotti and membrane dynamics during the late stages of NSC division. Selected frames from Video 1 are shown. Time in minutes relative to the start of imaging is shown, along with the diameter of the cytokinetic pore in microns. To the right, a kymograph of Pavarotti and membrane dynamics across the cytokinetic pore is shown.

The ICB forms when the central spindle becomes compacted in the cytokinetic pore through the action of the first furrowing step (Fig. 1A). The region of the central spindle with overlapping microtubules coated with proteins such as PRC1 and centralspindlin form the midbody^14,15^ while the adjacent flanking arms are the sites of constriction^5,16^. Flanking arm constriction gives rise to the characteristic ICB shape with two thin tubes surrounding a thicker midbody^3,11–13,17^. In cultured cells, membrane tension generated by migration of the nascent sibling cells contributes to the structure of the ICB and is also thought to drive arm constriction. While tension appears to be necessary for the formation and function of the ICB, abscission requires low tension. These different requirements suggests that cells must carefully regulate membrane tension to complete the final steps of cytokinesis. However, cells within tissues face mechanical constraints that could prevent tension generation by migration-based mechanisms. To understand ICB constriction in such an environment, we examined the asymmetrically dividing neural stem cells (NSCs) of the *Drosophila* larval brain (Fig. 1B).

NSCs divide rapidly, generating a larger self-renewed stem cell and a smaller neural precursor^18–20^. The initial furrow formation in NSCs is well-characterized, with both spindle- and polarity-dependent signals positioning the actomyosin ring^21–26^. As the initial furrow completes the first phase of NSC furrowing, cortical actomyosin flows are directed away from the equatorial region^27,28^. However, the subsequent ICB phase is poorly understood, including its structure and requirements for constriction. Here we investigate the mechanisms controlling this critical intermediate phase of cytokinesis in tissue-resident stem cells.

## Results

### The NSC intercellular bridge is a constricting midbody that lacks conventional flanking arms

Larval brain NSCs divide in a highly constrained environment, surrounded on one side by a cortex glial cell and the other by progeny from previous asymmetric divisions (Fig. 1B). We sought to understand how the NSC ICB functions in this environment given the extended structure of canonical ICBs. ICB arms are membranous tubes filled with microtubules that lack midbody proteins such as centralspindlin and PRC1 (Fascetto in *Drosophila*)^14,15,29^. We examined the structure of the NSC ICB using super resolution microscopy while monitoring the plasma membrane with PLCδ-PH along with microtubules or the centralspindlin protein MKLP1 (Pavarotti in *Drosophila*). The ICB forms when the cytokinetic pore reaches a diameter of approximately 1.5 μm when microtubules, Pavarotti (Pav), and other proteins become compacted within the pore. We observed several important differences between NSC and canonical ICBs (Fig. 1C-H; Video 1). First, the NSC ICB appeared to consist solely of the midbody, as no part of the membrane tube lacked Pav (Fig. 1H; Video 1). Second, the NSC ICB constricted at the midbody (Fig. 1H; Video 1). Finally, while formation of the canonical ICB is correlated with migration of nascent sibling cell bodies away from one another, the NSC nascent sibling membranes move in the opposite direction, becoming compressed against one another (Fig. 1E). Together, these data suggest that asymmetrically dividing neuroblasts form an intercellular bridge that lacks the conventional tripartite organization, instead consisting solely of a midbody.

### Sheets of plasma membrane oscillate near the cytokinetic pore during ICB formation and constriction

The plasma membrane is remodeled near the cytokinetic pore late in cytokinesis^30,31^. We examined the membrane remodeling in detail to determine if it is related to ICB formation or constriction. We imaged the full volume of the cytokinetic pore and surrounding membrane along with microtubules, revealing a highly dynamic process where sheets of plasma membrane formed simultaneously with midbody formation (Fig. 2A; Video 2). The sheets are formed in an oscillatory pattern such that they appear as waves emanating from the pore that extended into the cytoplasm (Fig. 2A,B; Video 2). The waves appeared predominantly on the side of the pore of the larger, NSC nascent sibling, although we cannot exclude the possibility that some membrane remodeling occurs on the other side. The oscillations of plasma membrane sheets continued through the ICB constriction process. Thus, we conclude that ICB formation and constriction is correlated with dramatic remodeling of the plasma membrane around the pore, where large sheets of oscillating membrane form around the pore for the duration of ICB thinning.

**Figure 2:**
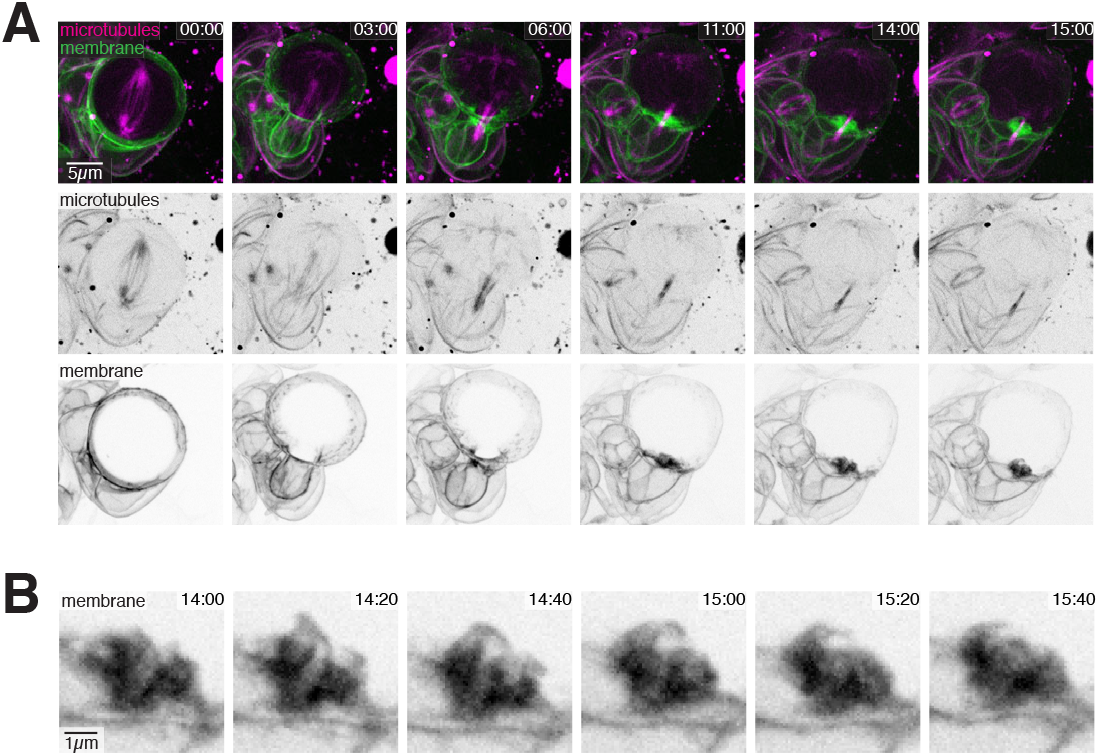
Plasma membrane oscillations near the cytokinetic pore during ICB formation and constriction. A) Membrane and microtubule dynamics during the late stages of NSC asymmetric division. Selected frames from Video 2 are shown. Maximum intensity projections of multiple optical sections spanning the cleavage furrow are shown. Time relative to anaphase onset is indicated. B) An example membrane oscillation. The cytokinetic pore and surrounding membrane is shown over the course of a single membrane oscillation. Time relative to anaphase onset is indicated.

### Plasma membrane oscillations near the pore are driven by F-actin

We sought to understand the cellular processes that drive membrane remodeling near the ICB pore. We imaged F-actin along with the membrane to determine if actin polymerization might be correlated with the oscillating membrane sheets as F-actin has been reported to be localized to protrusions that form late in NSC cytokinesis^31^. When imaging F-actin, we observed dense actin networks surrounding the membrane sheets, and these networks were highly correlated with sheet oscillations (Fig. 3A,B; Video 3). While a small amount of cortical actin was present before membrane sheet formation, a large burst of actin appeared when the deformations of the membrane around the pore began. Clouds of actin that surrounded the membrane sheets continued during the ICB thinning process. While strong F-actin signal surrounded the membrane oscillations near the pore, the signal within the pore was much more limited, especially early in the process when the pore was at its largest.

**Figure 3:**
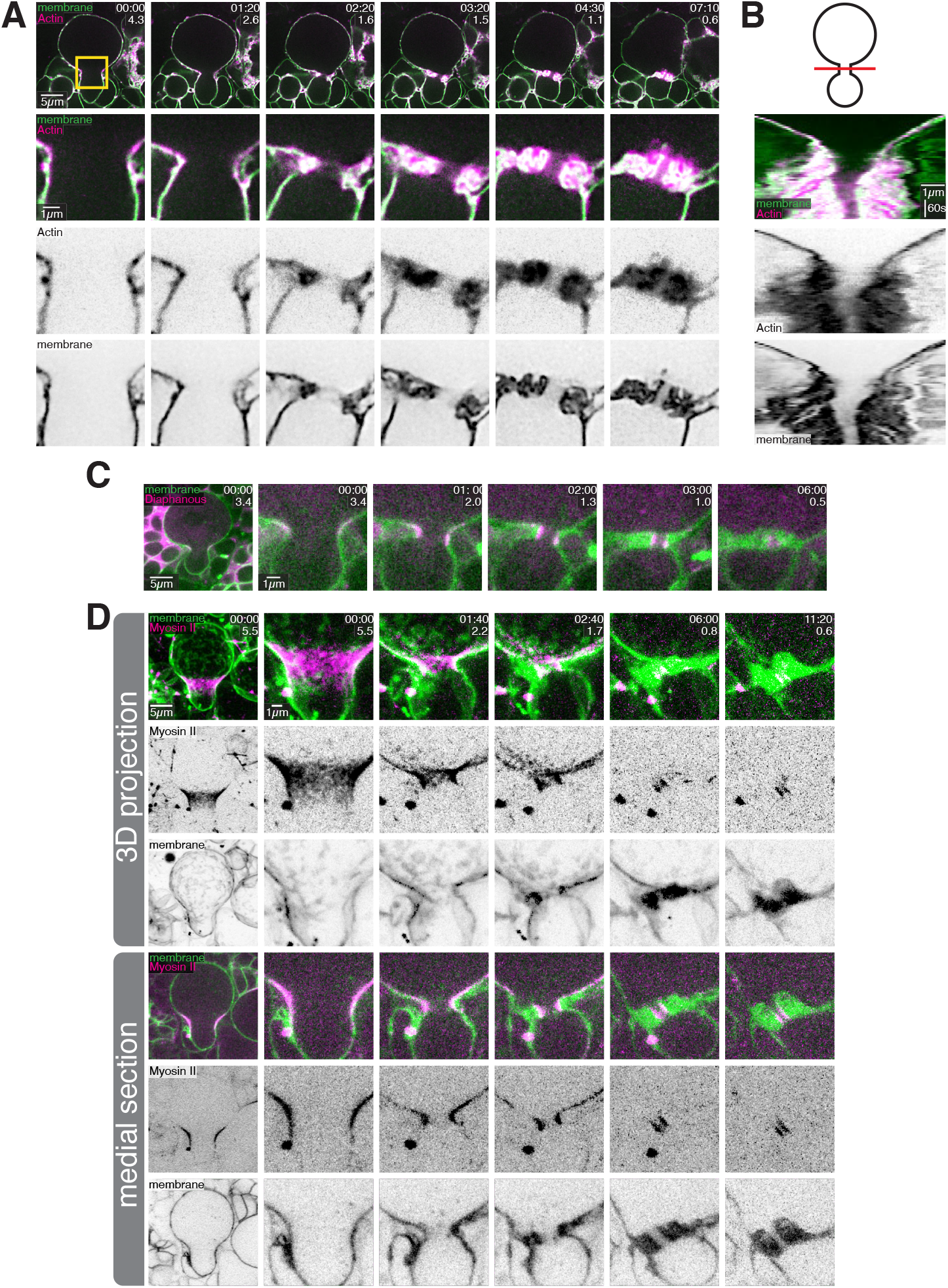
F-actin localizes to plasma membrane waves near the cytokinetic pore. A) F-actin and membrane dynamics during the late stages of NSC division. Selected frames from Video 3 are shown. The top row depicts a medial section where the entire cell is visible. The rows below are a zoomed-in view of the cleavage furrow. Time relative to the first frame is indicated, along with the diameter of the cytokinetic pore in microns. Image deconvolution was applied. B) A kymograph of F-actin and membrane dynamics across the cytokinetic pore. C) Diaphanous and membrane dynamics during the late stages of NSC asymmetric division. Selected frames are shown from Video 3. A medial section where the entire cell is visible (left) is next to a timelapse of a zoomed-in view of the cleavage furrow. Time relative to the first frame is indicated, along with the diameter of the cytokinetic pore in microns. D) Myosin II and membrane dynamics during the late stages of NSC asymmetric division. Selected frames are shown from Video 3. The top 3 rows depicts maximum intensity projections of an entire hemisphere of the dividing NSC. The bottom row is a medial section. The left column shows a view of the whole cell, and the columns to the right are zoomed-in views of the cleavage furrow. Time relative to the first frame is indicated, along with the diameter of the cytokinetic pore in microns.

We sought to identify the cytoskeletal proteins that regulate or cooperate with sheet-associated actin. The formin F-actin nucleator Diaphanous (Dia), was strongly enriched on the wall of the ICB (Fig. 3C; Video 3), but we did not detect it in the surrounding membrane sheets. We observed Myosin II broadly distributed on the equatorial cortex during the initial furrowing stage (Fig 3D; Video 3). Immediately before ICB formation, Myosin II dissipated from the furrow as previously described^28^, before becoming strongly enriched on the ICB wall. We did not detect any significant Myosin II localization with the membrane sheets that surround the ICB pore. We conclude that the F-actin associated with the membrane sheets surrounding the ICB pore is not nucleated by Dia and is not part of an actomyosin contractile network.

### Arp2/3 actin filament nucleation is required for NSC ICB thinning

We also examined the localization of a nucleator of branched actin networks, the Arp2/3 complex. The Arp-3 subunit localized specifically to sites of plasma membrane remodeling surrounding the pore and was recruited shortly before the burst of actin that coincides with membrane sheet formation (Fig. 4A; Video 4). The Arp2/3 activator SCAR (aka WAVE complex) localizes to the NSC furrow^31^ and we observed localization to the membrane sheets that form around the ICB (Fig. 4B; Video 4). To examine the function of the Arp2/3 complex in membrane remodeling and ICB function, we acutely inhibited its activity using the chemical inhibitor CK-666. Arp2/3 inhibition before ICB formation completely abrogated membrane sheet formation and caused stalling of the furrow at a diameter of approximately 1.5 μm (Fig. 4C,D; Video 4). We also inhibited Arp2/3 after membrane sheet oscillations had initiated and the midbody had formed. In these cells, membrane sheet dynamics and ICB constriction abruptly stopped (Fig. 4E; Video 4). We conclude that Arp2/3 is required for the membrane oscillations near the ICB, and for ICB formation and constriction.

**Figure 4:**
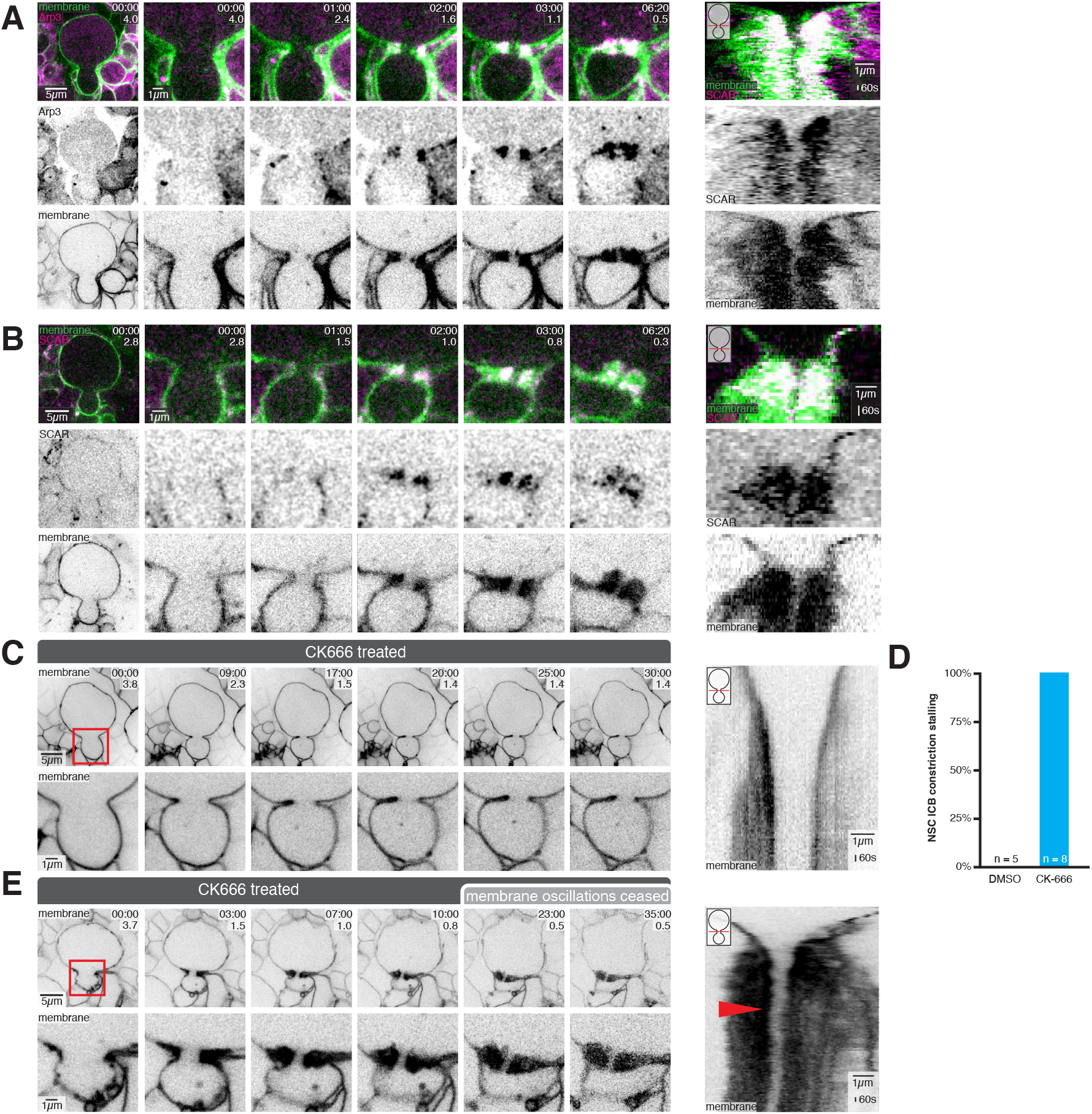
Arp2/3 is required for plasma membrane oscillations and ICB constriction. A) Arp2/3 complex and membrane dynamics during the late stages of NSC division. Selected frames from Video 4 are shown. A medial section where the entire cell is visible (left) is next to a timelapse of a zoomed-in view of the cleavage furrow. Time relative to the first frame is indicated, along with the diameter of the cytokinetic pore in microns. To the right is a kymograph of Arp2/3 complex and membrane dynamics across the cytokinetic pore. B) SCAR complex and membrane dynamics during the late stages of NSC division. Selected frames from Video 4 are shown. A medial section where the entire cell is visible (left) is next to a timelapse of a zoomed-in view of the cleavage furrow. Time relative to the first frame is indicated, along with the diameter of the cytokinetic pore in microns. To the right is a kymograph of SCAR complex and membrane dynamics across the cytokinetic pore. C) The effects of Arp2/3 inhibition on membrane dynamics during the late stages of NSC division. Selected frames from Video 4 are shown. The NSC was treated with the Arp2/3 inhibitor CK666 before the formation of the intercellular bridge (ICB). The left column depicts a medial section where the entire cell is visible. The bottom row is a zoomed-in view of the cleavage furrow. Time relative to start of imaging is indicated, along with the diameter of the cytokinetic pore in microns. To the right is a kymograph of membrane dynamics across the cytokinetic pore for the CK666-treated NSC. D) Quantification of the effect of inhibition of Arp2/3 complex before ICB formation on late cytokinesis. NSCs from larval brains incubated with DMSO (carrier) or CK-666 before ICB formation were scored for stalled ICB constriction where the ICB did not reach a pore diameter of approximately 200 nm. E) The same as in (C), except the NSC was treated with the Arp2/3 inhibitor CK666 after the formation of the intercellular bridge (ICB) and membrane oscillations had started. To the right is a kymograph of membrane dynamics across the cytokinetic pore for the CK666-treated NSC. The red arrow approximates when Arp2/3 was inhibited.

## Discussion

In this study, we addressed a fundamental gap in our understanding of cytokinesis - how cells complete division when constrained within tissues, where they may not be able to generate tension across the ICB through migration like isolated cells do. We make the key discovery that Drosophila larval brain NSCs utilize a distinct ICB structure during cytokinesis, characterized by constriction focused at the central midbody rather than flanking arms. We also reveal previously unrecognized oscillatory membrane dynamics surrounding the ICB that depend on Arp2/3-mediated branched actin networks.

We propose that the unique structure of the NSC ICB, lacking the conventional tripartite organization with flanking arms, may represent an adaptation for cells dividing within tissues. While isolated cells can generate tension across their ICB through migration, tissue-resident cells face mechanical constraints that may prevent such movement. Indeed, we observe that NSC daughter cells become compressed against each other rather than separating when the ICB forms (Fig. 1E). The focused constriction at the midbody, rather than along extended arms, could allow these cells to complete cytokinesis without requiring active migration of the nascent sibling cells away from one another.

The dramatic membrane remodeling we observe around the ICB, with oscillating membrane sheets driven by Arp2/3-dependent actin networks, may serve to generate localized forces that facilitate bridge constriction in the absence of migration-based tension. This is reminiscent of how Arp2/3 remodels membranes during processes like endocytosis, where branched actin networks provide force for membrane deformation^32,33^. The correlation between membrane oscillations and ICB constriction, along with the complete block of constriction when Arp2/3 is inhibited, suggests these dynamics play an essential role. We propose that the membrane waves could generate mechanical forces that aid in bridge closure or help regulate membrane tension across the ICB in a way that promotes sufficient constriction to allow abscission to take place.

## Resource Availability

### Lead Contact

Contact the Lead Contact, Kenneth Prehoda, for further information or to request resources and reagents.

### Materials Availability

No new reagents were generated in this study.

### Data and Code Availability

Raw data available from the corresponding author on request.

### Experimental Model and Subject Details

#### Fly Strains

A Worniu-GAL4 driver line was used to drive tissue specific expression of UAS controlled transgenes in neural stem cells (NSCs). Membrane dynamics were imaged using various membrane markers. UAS-PLCδ-PH-GFP and UAS-PLCδ-PH-mCherry express the pleckstrin homology domain of human PLCδ tagged with GFP or mCherry, and binds to the plasma membrane lipid phosphoinositide PI(4,5)P2. UAS-GRP1-PH-GFP expresses the pleckstrin homology domain of GRP1, and binds to the plasma membrane lipid phosphoinositide PI(3,4,5)P3 34. UAS-Farnesyl-GFP expresses the C-terminal region of human K-Ras tagged with GFP which becomes farnesylated and membrane-anchored in cells. F-Actin was visualized using UAS-GMA-GFP, which expresses a GFP tagged actin binding domain of Moesin, and UAS-Lifeact-mRuby. Microtubules were imaged using UAS-Zeus-mCherry. Anillin was imaged using UAS-Anillin-GFP. Arp2/3 was imaged using UAS-Arp3-GFP. Centralspindlin dynamics were imaged using GFP tagged Pavarotti (Pav) protein under control of ubiquitin regulatory sequences. The formin, Diaphanous, was imaged using UAS-Diaphanous-GFP. The SCAR/WAVE complex was imaged using Sra1/Cyfip endogenously tagged with eGFP using CRISPR/Cas935. Myosin II was imaged using GFP tagged Spaghetti squash, the regulatory light chain of non-muscle type II Myosin, expressed from its endogenous promoter36.

### Method Details

#### Live Imaging

To obtain brain explants, third instar *Drosophila* larvae were dissected in Schneider’s Insect Media (SIM) to isolate the central nervous system. Brain explants were mounted on sterile poly-D-lysine coated 35mm glass bottom dish (ibidi Cat#81156) containing modified minimal hemolymph-like solution (HL3.1). Brain explants were imaged using a Nikon Eclipse Ti-2 Yokogawa CSU-W1 SoRa spinning disk microscope equipped dual Photometrics Prime BSI sCMOS cameras using a 60x H2O objective. 488 nm light was used to illuminate GFP tagged proteins and 561 nm light was used to illuminate mCherry and mRuby tagged proteins. Super resolution imaging was achieved by using SoRa (super resolution through optical photon reassignment) optics 37. NSCs were identified by their large size, location in the central nervous system, and the use of NSC specific tissue driver lines. Time lapse imaging of cleavage furrow and intercellular bridge dynamics was achieved by refocusing the imaging plane on the medial plane of the cleavage furrow and intercellular bridge, along the apical-basal axis just before capturing each frame. Pharmacological inhibition of the Arp2/3 complex was performed using 2 mM CK666 solubilized in DMSO.

#### Image Processing and Analysis

Imaging data was processed using ImageJ (FIJI package). For some movies, the bleach correction tool was used to correct for photobleaching. To reduce noise in Arp3-GFP and SCAR-GFP images, Gaussian blur was applied. When image deconvolution was applied, deconvolution was performed using Nikon Elements standard 2D deconvolution mode and noted in the figure/video legend. For cytokinetic pore size measurements, medial sections were used to measure the width of the cytokinetic pore.

Quantifying the effects of Arp2/3 inhibition of membrane dynamics during the late stages of NSC division: Closure of the cytokinetic pore was measured for CK666-treated NSCs and compared to untreated NSCs. For quantifying the dynamics of the cytokinetic pore size, medial sections (along the apical-basal axis) were used to measure the width of the cytokinetic pore.

### Key Resources Table

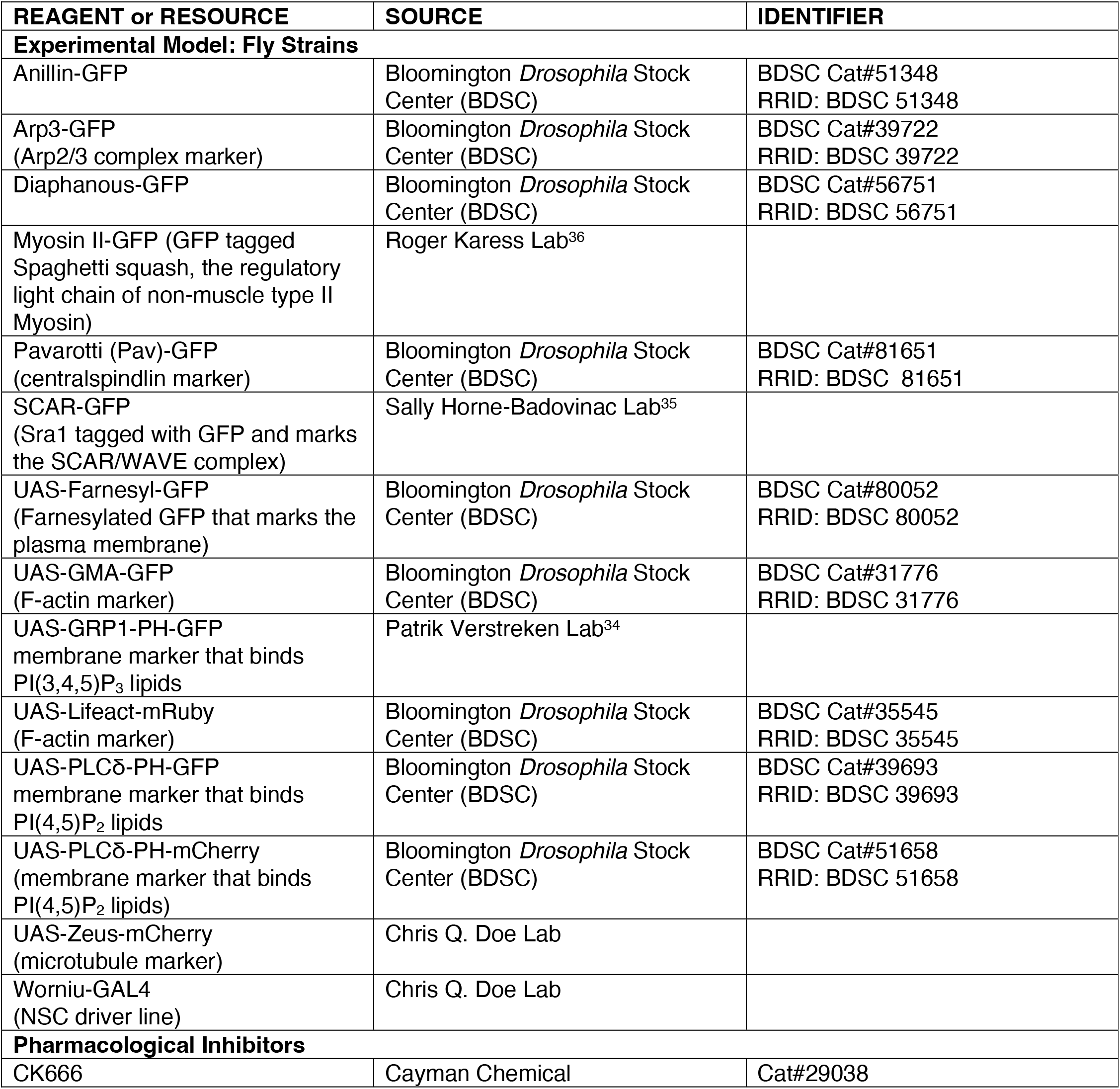

## Video Legends

**Video 1: Larval brain neural stem cell intercellular bridges are midbodies without flanking arms**

Part 1: Membrane and microtubule dynamics during the late stages of NSC division. Super resolution videos of an NSC expressing Zeus-mCherry “Microtubules” and the membrane marker Farnesyl-GFP “membrane”. The top row depicts a medial section where the entire cell is visible. The bottom row is a zoomed-in view of the cleavage furrow. Time relative to start of imaging is indicated.

Part 2: Phosphoinositide localization during the late stages of NSC division. Super resolution videos of an NSC expressing the membrane markers UAS-GRP1-GFP “PI(3,4,5)P3” and UAS-PLCδ-PH-mCherry “PI(4,5)P2”. The top row a medial section where the entire cell is visible. The bottom row is a zoomed-in view of the cleavage furrow. Time relative to start of imaging is indicated.

Part 3: Anillin and membrane dynamics during the late stages of NSC division. Super resolution videos of an NSC expressing Anillin-GFP “Anillin” and the membrane marker UAS-PLCδ-PH-mCherry “membrane”. The top row depicts a medial section where the entire cell is visible. The bottom row is a zoomed-in view of the cleavage furrow. Time relative to start of imaging is indicated.

Part 4: Pavrotti and membrane dynamics during the late stages of NSC division. Super resolution videos of an NSC expressing Pavarotti-GFP “Pavarotti” and the membrane marker UAS-PLCδ-PH-mCherry “membrane”. The top row depicts a medial section where the entire cell is visible. The bottom row is a zoomed-in view of the cleavage furrow. Time relative to start of imaging is indicated.

**Video 2: Plasma membrane oscillations near the cytokinetic pore during intercellular bridge formation and constriction**

Part 1: 3-dimensional view of membrane and microtubule dynamics during the late stages of NSC division. Super resolution videos of an NSC expressing Zeus-mCherry “Microtubules” and the membrane marker Farnesyl-GFP “membrane”. Maximum intensity projections of multiple optical sections spanning the cleavage furrow are shown. Time relative to anaphase onset is indicated.

Part 2: Top-down view Anillin and membrane dynamics during the late stages of NSC division. Super resolution videos of an NSC expressing Anillin-GFP “Anillin” and the membrane marker UAS-PLCδ-PH-mCherry “membrane”. A single optical section is shown. Time relative to start of cytokinetic pore constriction is indicated.

**Video 3: F-actin localizes to plasma membrane waves near the cytokinetic pore**

Part 1: Actin and membrane dynamics during the late stages of NSC division. Super resolution videos of an NSC expressing Lifeact-mRuby “Actin” and the membrane marker UAS-PLCδ-PH-GFP “membrane”. The top row depicts a medial section where the entire cell is visible. The bottom row is a zoomed-in view of the cleavage furrow. Time relative to anaphase onset is indicated. Image deconvolution was applied.

Part 2: Top-down view Actin and membrane dynamics during the late stages of NSC division. Super resolution videos of an NSC expressing GMA-GFP “Actin” and the membrane marker UAS-PLCδ-PH-mCherry “membrane”. A single optical section is shown. Time relative to start of cytokinetic pore constriction is indicated.

Part 3: Myosin II and membrane dynamics during the late stages of NSC division. Super resolution videos of an NSC expressing Myosin II-GFP “Myosin II” and the membrane marker UAS-PLCδ-PH-mCherry “membrane”. The top row depicts maximum intensity projections of an entire hemisphere of the dividing NSC. The bottom row is a medial section. The left column shows a view of the whole cell, and the columns to the right are zoomed-in views of the cleavage furrow. Time relative to anaphase onset is indicated.

Part 4: Diaphanous and membrane dynamics during the late stages of NSC division. Super resolution videos of an NSC expressing Diaphanous-GFP “Diaphanous” and the membrane marker UAS-PLCδ-PH-mCherry “membrane”. The top row depicts a medial section where the entire cell is visible. The bottom row is a zoomed-in view of the cleavage furrow. Time relative to start of imaging is indicated.

**Video 4: Arp2/3 is required for plasma membrane oscillations and intercellular bridge constriction**

Part 1: Arp2/3 complex and membrane dynamics during the late stages of NSC division. Super resolution videos of an NSC expressing Arp3-GFP “Arp 3” and the membrane marker UAS-PLCδ-PH-mCherry “membrane”. The top row depicts a medial section where the entire cell is visible. The bottom row is a zoomed-in view of the cleavage furrow. Time relative to start of imaging is indicated. In post-processing, Guassian blur was applied to the GFP channel to reduce noise.

Part 2: SCAR/WAVE complex and membrane dynamics during the late stages of NSC division. Super resolution videos of an NSC expressing SCAR-GFP “SCAR” and the membrane marker UAS-PLCδ-PH-mCherry “membrane”. The top row depicts a medial section where the entire cell is visible. The bottom row is a zoomed-in view of the cleavage furrow. Time relative to start of imaging is indicated. In post-processing, Guassian blur was applied to the GFP channel to reduce noise.

Part 3: The effects of Arp2/3 inhibition on membrane dynamics during the late stages of NSC division. Super resolution videos of NSCs expressing membrane marker UAS-PLCδ-PH-GFP “membrane” and treated with the Arp2/3 inhibitor CK666. The left column depicts a medial section where the entire cell is visible. The bottom row is a zoomed-in view of the cleavage furrow. Time relative to start of imaging is indicated. Arp2/3 inhibition occurs early on during cytokinesis in the first cell, whereas Arp2/3 becomes inhibited later in the second cell.

## Acknowledgments

We thank Adam Fries for maintaining the microscope used in this study. We thank Sally Horne-Badovinac for the SCAR-GFP fly line. We thank Patrik Verstreken for the UAS-GRP1-PH-GFP fly line. We thank Chris Q. Doe for the Worniu-GAL4 and UAS-Zeus-mCherry fly lines. This work was supported by NIH grants R35GM127092 and K99GM147601.

## Author Contributions

B.L. and K.E.P. designed the experiments. B.L. performed the experiments. B.L. and K.E.P analyzed the data, prepared the figures, and wrote the manuscript.

